# Evolution of the open-access CIViC knowledgebase is driven by the needs of the cancer variant interpretation community

**DOI:** 10.1101/2021.06.13.448171

**Authors:** Kilannin Krysiak, Arpad M Danos, Susanna Kiwala, Joshua F McMichael, Adam C Coffman, Erica K Barnell, Lana Sheta, Jason Saliba, Cameron J Grisdale, Lynzey Kujan, Shahil Pema, Jake Lever, Nicholas C Spies, Andreea Chiorean, Damian T Rieke, Kaitlin A Clark, Payal Jani, Hideaki Takahashi, Peter Horak, Deborah I Ritter, Xin Zhou, Benjamin J Ainscough, Sean Delong, Mario Lamping, Alex R Marr, Brian V Li, Wan-Hsin Lin, Panieh Terraf, Yasser Salama, Katie Campbell, Kirsten M Farncombe, Jianling Ji, Xiaonan Zhao, Xinjie Xu, Rashmi Kanagal-Shamanna, Kelsy C Cotto, Zachary L Skidmore, Jason R Walker, Jinghui Zhang, Aleksandar Milosavljevic, Ronak Y Patel, Rachel H Giles, Raymond H Kim, Lynn M Schriml, Elaine R Mardis, Steven JM Jones, Gordana Raca, Shruti Rao, Subha Madhavan, Alex H Wagner, Obi L Griffith, Malachi Griffith

## Abstract

CIViC (Clinical Interpretation of Variants in Cancer; civicdb.org) is a crowd-sourced, public domain knowledgebase composed of literature-derived evidence characterizing the clinical utility of cancer variants. As clinical sequencing becomes more prevalent in cancer management, the need for cancer variant interpretation has grown beyond the capability of any single institution. With nearly 300 contributors, CIViC contains peer-reviewed, published literature curated and expert-moderated into structured data units (*Evidence Items*) that can be accessed globally and in real time, reducing barriers to clinical variant knowledge sharing. We have extended CIViC’s functionality to support emergent variant interpretation guidelines, increase interoperability with other variant resources, and promote widespread dissemination of structured curated data. To support the full breadth of variant interpretation from basic to translational, including integration of somatic and germline variant knowledge and inference of drug response, we have enabled curation of three new evidence types (predisposing, oncogenic and functional). The growing CIViC knowledgebase distributes clinically-relevant cancer variant data currently representing >2500 variants in >400 genes from >2800 publications.

## Introduction

CIViC is an open-access, open-source knowledgebase for the expert crowdsourcing of Clinical Interpretation of Variants in Cancer (civicdb.org)^1,2^. The critical distinguishing features of the CIViC initiative, in contrast to other somatic cancer variant resources, stem from its strong commitment to open-access and data provenance. Six founding principles to maintain a *freely* and *computationally accessible* resource with *transparency*, an *open license*, and *interdisciplinary* participation to support *community consensus* underlie this resource. From this foundation, we hypothesized that CIViC could strategically apply crowdsourcing techniques to achieve widespread adoption. Curators extract detailed evidence of the clinical significance of variants in cancer from the peer-reviewed, published literature to contribute to the resource. Crowdsourced contributions are moderated after submission by expert Editors, individuals who are familiar with CIViC standard operating procedures, have undergone training, and have field-relevant expertise (e.g., graduate/medical degrees) in a gatekeeper curation model^1,2^. CIViC was designed to encourage the development of community consensus by leveraging an interdisciplinary, international team of experts collaborating remotely within a centralized curation interface. Curated variant interpretations are made available through a web interface (no login required) and a well-documented, modern application programming interface (API), under a public domain (CC0) dedication. All software is available on GitHub under an MIT license. This approach has permitted global engagement with stakeholders in government, industry, and academia both nationally and internationally. Through community engagement, the capabilities of the CIViC knowledgebase and interface have been augmented to meet the needs of the field. The carefully crafted web interface, a performant API, and agile development process have made these improvements possible. We believe that the CIViC platform exemplifies successful implementation of a crowdsourced knowledgebase with clinical utility using a completely open model.

The existing annotation bottleneck associated with variant interpretation is well described^3^. The ever growing repertoire of variants associated with cancer has led to many variants falling outside of clinical guidelines, resulting in a need for resources to assist variant analysts, geneticists, and oncologists. For example, tyrosine kinase inhibitors are the standard of care in non-small cell lung cancers harboring canonical *EGFR* variants (e.g., p.T790M, p.L858R). However, many patients harbor non-canonical *EGFR* variants, which in one study represented 9% of *EGFR*-mutated patients, demonstrating the need for and potential clinical benefits of the aggregation of evidence for rare variants^4^. Additionally, many genes are known to contribute to tumorigenesis through variants of both germline and somatic origin (e.g., *TP53*, *VHL, BRCA1*), and the investigation of the clinical utility of somatic variants in germline interpretations, and vice versa, are increasingly being considered in clinical practice^5,6^. The validation of platforms that can quickly and effectively incorporate genomics to guide the diagnosis, prognosis, and treatment of cancers is required to alleviate existing bottlenecks within precision oncology.

Here, we describe the impact of the CIViC knowledgebase on the field through 1) engagement and contributions of external users, 2) adaptation to support emerging guidelines, 3) community-driven evolution of the data model and user interface, and 4) integration with external resources. Through this research, we exemplify how extensive and international adoption of the CIViC knowledgebase centralizes curation and improves the distribution of curated cancer variant evidence.

## Results

### Scaling up curation through community engagement

Since its initial release, the CIViC knowledgebase has undergone rapid expansion and adoption supporting a broader ecosystem of cancer variant resources. Due to the unrestricted licensing and open API, CIViC data consumers are not required to register their use; therefore, the complete picture of CIViC’s impact cannot be determined. However, established collaborations and self-identified data clients illustrate several types of integration and diverse stakeholders including academic, government and commercial entities as well as clinical and basic science resources (**Figure 1**)^7–10^. Most groups and individuals that interact with CIViC consume the data without creating new content. Web users span the globe (with more than 90,000 users from outside the United states) and visit, on average, 11 pages per session with the duration of such visits averaging ~7 minutes (**Figure 2**). However, anyone can create a login and contribute. Contributors who comment, submit new content, and suggest revisions to existing content are referred to as CIViC *Curators* (N=284 as of May 1st, 2021), the majority of which are individuals from outside of the Washington University in Saint Louis (WashU) community (only 40 of 284 active Curators have a WashU affiliation, the site of CIViC’s initial development) (**Figure 3a**). These Curators represent academic, governmental and commercial organizations^8,11^. Curator contributions take many forms and require varying degrees of effort, which supports curation activities from individuals with a wide range of interests and expertise (**Supplemental Figure 1**, **Table 1**). Among the more time-intensive activities is the curation and moderation of *Evidence Items*. As the foundational unit of CIViC, Evidence Items associate a variant with a clinical interpretation derived from published biomedical literature. Using Evidence curation as a proxy for community engagement, we observe external contributions currently exceed internal contributions (**Figure 3b**, **Supplemental Figure 2**). These data illustrate the success of CIViC in engaging hundreds of outside users in the knowledge curation process.

**Figure 1.**
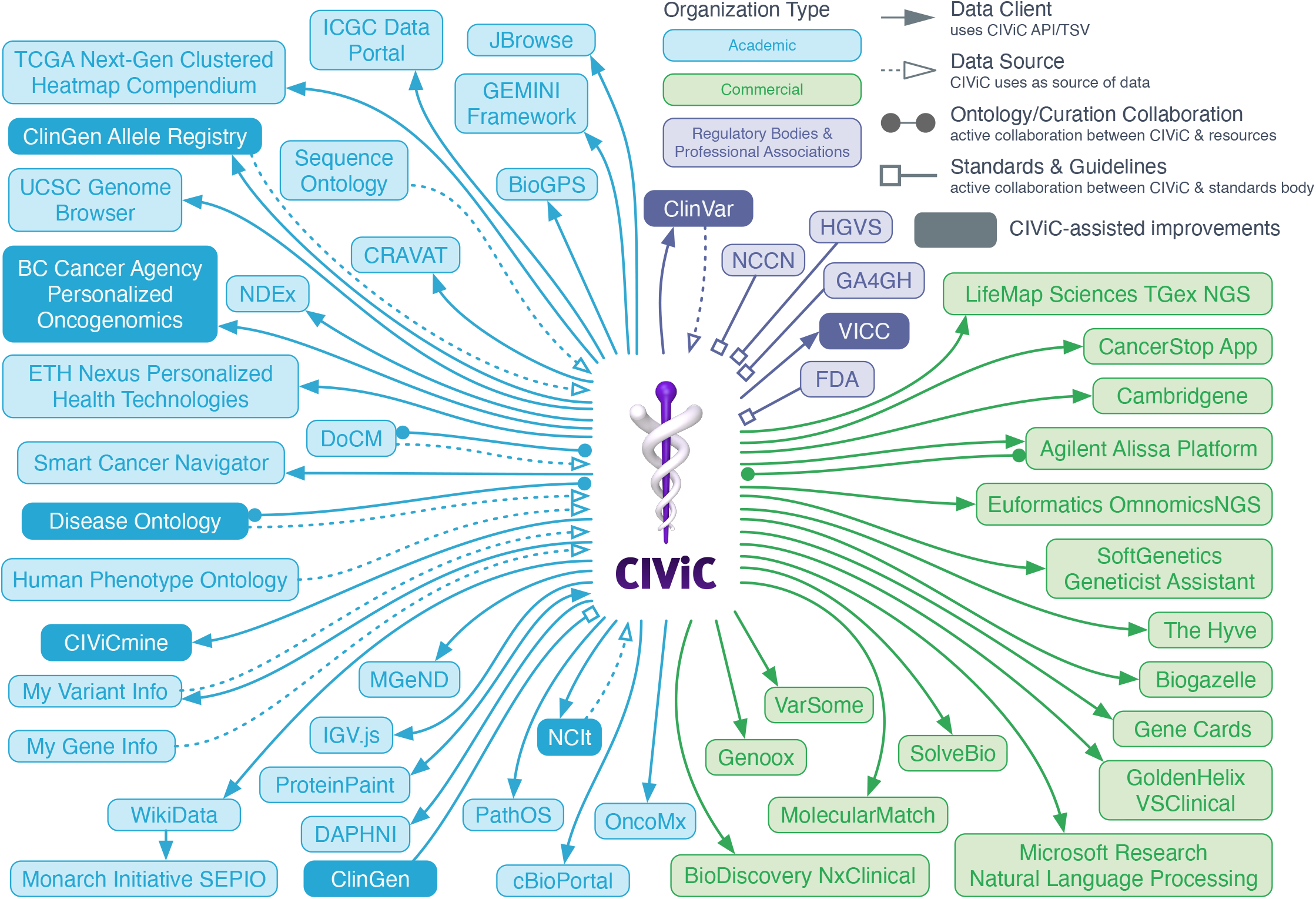
The CIViC Ecosystem. This web shows community engagement with the CIViC Interface. Colors represent the Organization Type (Regulatory / Professional Associations, Academic, and Commercial) and connection indicates the type of interaction with the organization.

**Figure 2.**
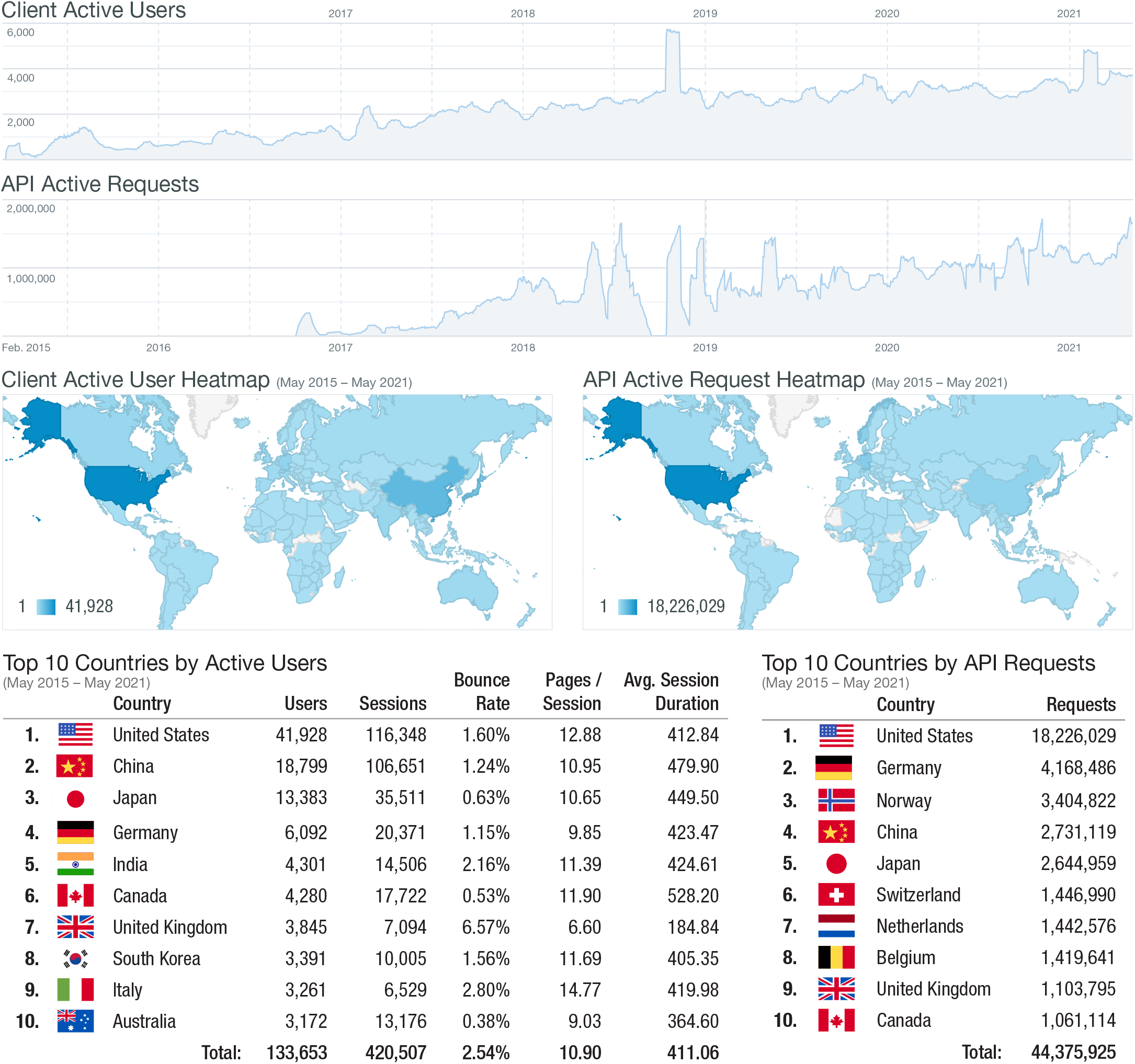
API and web usage statistics. Total engagements with the database are shown. The blue density plots show client users accessing CIViC through the web interface (top density plot) and pull requests from the CIViC API (bottom density plot). Heatmaps (middle) show the originating country (based on IP address) from users interacting with the web interface (left) and API (right). Activity from the top 10 countries is shown (bottom), demonstrating the number of users and individual sessions (new visits to the web interface). Users view approximately 11 pages per visit, spending on average 411 seconds in the web interface per visit.

**Figure 3.**
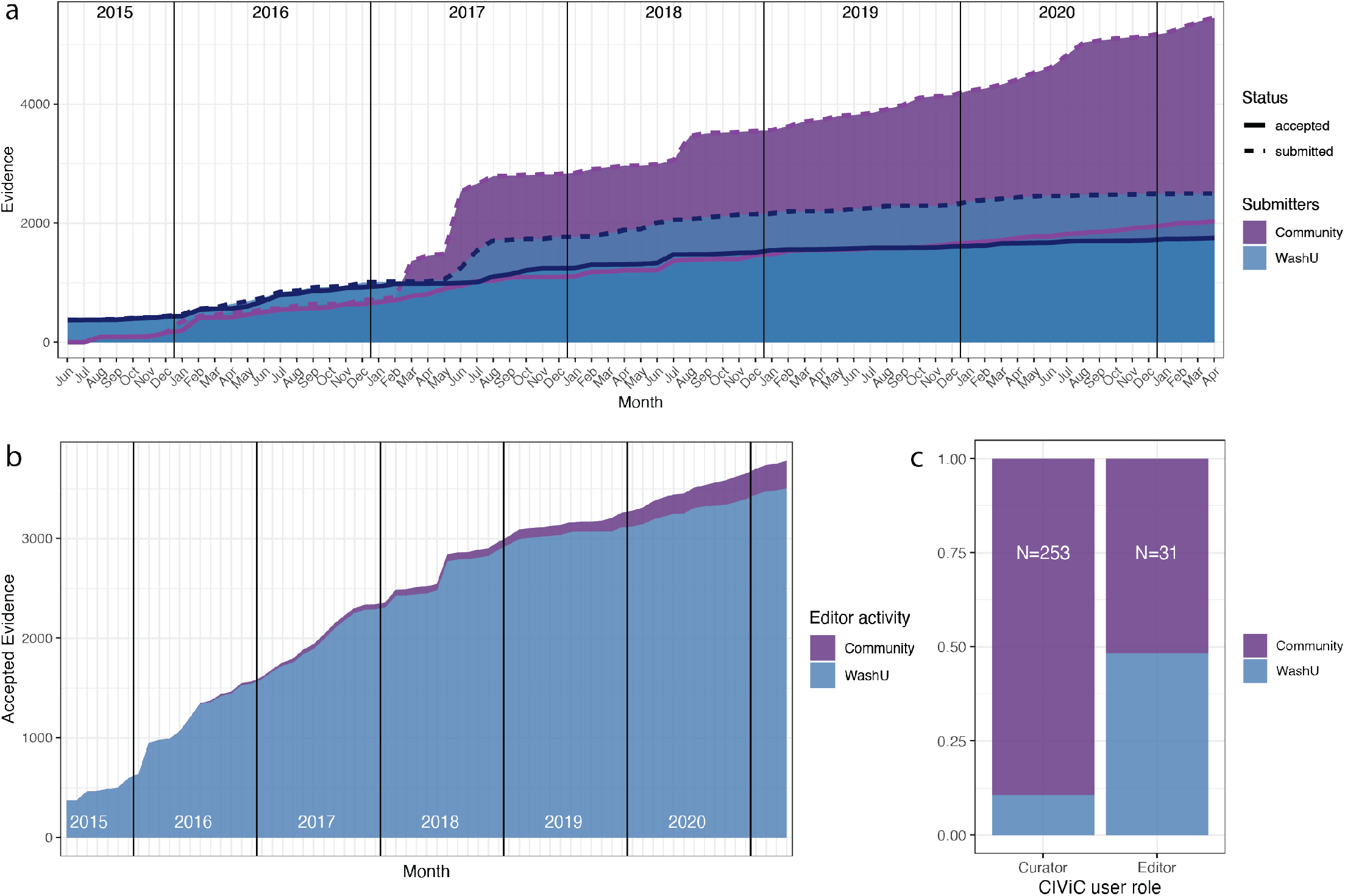
Moderation activity by contribution source. **a)** The current affiliations of Curators (only those who have performed curation activities, excluding Editors) and Editors working in the CIViC interface (Total = 284). **b)** At the inception of the knowledgebase, initial contributions to the database were performed by internal Curators (Washington University in St Louis, WashU). However, in 2017, external curation (Community) exceeded the internal contribution. To date, the gap between internal and external contribution continues to widen as new external users begin to adopt and contribute to the database. Curation activity has exceeded moderation activity of Editors, as represented by Accepted Evidence statements (solid lines) compared to Submitted Evidence Items (dashed lines). **c)** After the initial launch, external Curators were identified, trained and promoted to participate in moderation activities including accepting Evidence Items, as shown.

**Table 1.** CIViC curation statistics from original publication to current.

| Category | December 2016 | May 2021 | % Growth |
| --- | --- | --- | --- |
| Contributors | 58 | 284 | 490% |
| Total Evidence Items | 1703 | 7727 | 454% |
| Variants | 731 | 2621 | 359% |
| Publications | 1076 | 2895 | 269% |
| Total Accepted Evidence Items | 1678 | 3783 | 225% |
| Drugs | 291 | 463 | 159% |
| Genes | 283 | 442 | 156% |
| Diseases | 209 | 304 | 145% |

A subset of expert Curators (CIViC *Editors*; N=31) are selected and trained to moderate submitted content and maintain the quality of the data in the resource, as previously described^1,2^. External evidence submissions are essential to the scalability and overall success of CIViC. However, increased curation has led to the accumulation of content that must be moderated by expert Editors. These submissions can be viewed in the interface with warnings that they have not yet been moderated by Editors. The rapid influx of Evidence Item curation has overtaken Editor capacity, resulting in a large amount of evidence in a pending state (**Table 1, Figure 3b**). This was expected given the expertise and commitment required of Editors volunteering their time. Having anticipated the moderation bottleneck, we developed training programs to improve the quality of Curator submissions and expedite the promotion of qualified individuals into Editor roles. We published a standard operating procedure (SOP) for CIViC curation efforts^2^, generated training videos and tutorials, and developed extensive help documentation, available at docs.civicdb.org. Transparency of curation and moderation was improved through the addition of conflict of interest statements, which are required for all Editors at least annually. The success of CIViC has enabled engagement with Editors from collaborating organizations. For example, members of the Personalized OncoGenomics program^12^ (NCT02155621) at BC Cancer (British Columbia, Canada) have been trained as Editors, allowing them to curate and moderate CIViC evidence associated with real-world precision oncology cases, while also providing feedback to improve CIViC integration within their variant interpretation workflow. Since the inception of CIViC, 3506 EIDs have been accepted by 15 WashU-affiliated Editors, and 277 by 16 community Editors (7.3%). While a more recent snapshot of Editor activity (2020-2021) shows 25% (130/520) of EIDs have been accepted by community Editors (**Figure 3c**). CIViC continues to produce educational material in various formats to support high-quality curation and engage more community Curators interested in becoming Editors. These efforts are further enhanced by our extensive collaboration with the Clinical Genome Resource (ClinGen) Somatic Cancer Clinical Domain Working Group (SC-CDWG; https://clinicalgenome.org/curation-activities/somatic/)^13^. To date, 4/16 community Editors were recruited from the SC-CDWG, with 7 additional Editors currently in training. Through this collaboration, CIViC has become the variant curation platform for current and future ClinGen Somatic Cancer Variant Curation Expert Panels (SC-VCEPs).

### Guideline-driven evolution of CIViC variant tiering and classification

Several organizations have published guidelines for evaluating, interpreting, reporting and cataloging evidence pertaining to cancer variants and their structured representation in databases^14–16^. The CIViC team engaged members of those organizations and the community to determine how to best integrate these guidelines into curation efforts. This engagement led to modifications of the CIViC knowledgebase schema and incorporation of new features that support the most current recommendations^7,17^.

An example of CIViC’s responsiveness to emergent recommendations is our incorporation of the widely accepted 2017 Association for Molecular Pathology, American Society of Clinical Oncology, and College of American Pathologists (AMP/ASCO/CAP) guidelines for the interpretation and reporting of sequence variants in cancers^14^. To facilitate curation in a manner compliant with the AMP/ASCO/CAP recommendations, we developed a new entity: the CIViC *Assertion* (**Figure 4**). CIViC Assertions aggregate Evidence Items for a given variant-disease or variant-disease-therapy combination to provide an overarching “state of the field” clinical significance classification. Assertions support different structured data fields than Evidence Items, including AMP/ASCO/CAP tier and level classifications and associations with National Comprehensive Cancer Network (NCCN) guidelines and Food and Drug Administration (FDA) approvals. Many clinical labs and resources share final variant classifications without detailing the underlying evidence used to make these classifications. In contrast, CIViC Assertions are visible to the public while they are being created, clearly labeled as not yet complete, and open to community feedback and revision. Therefore, the developing Assertion and underlying evidence can be assessed by the public at all steps of the process, prior to formal, finalized variant classifications.

**Figure 4.**
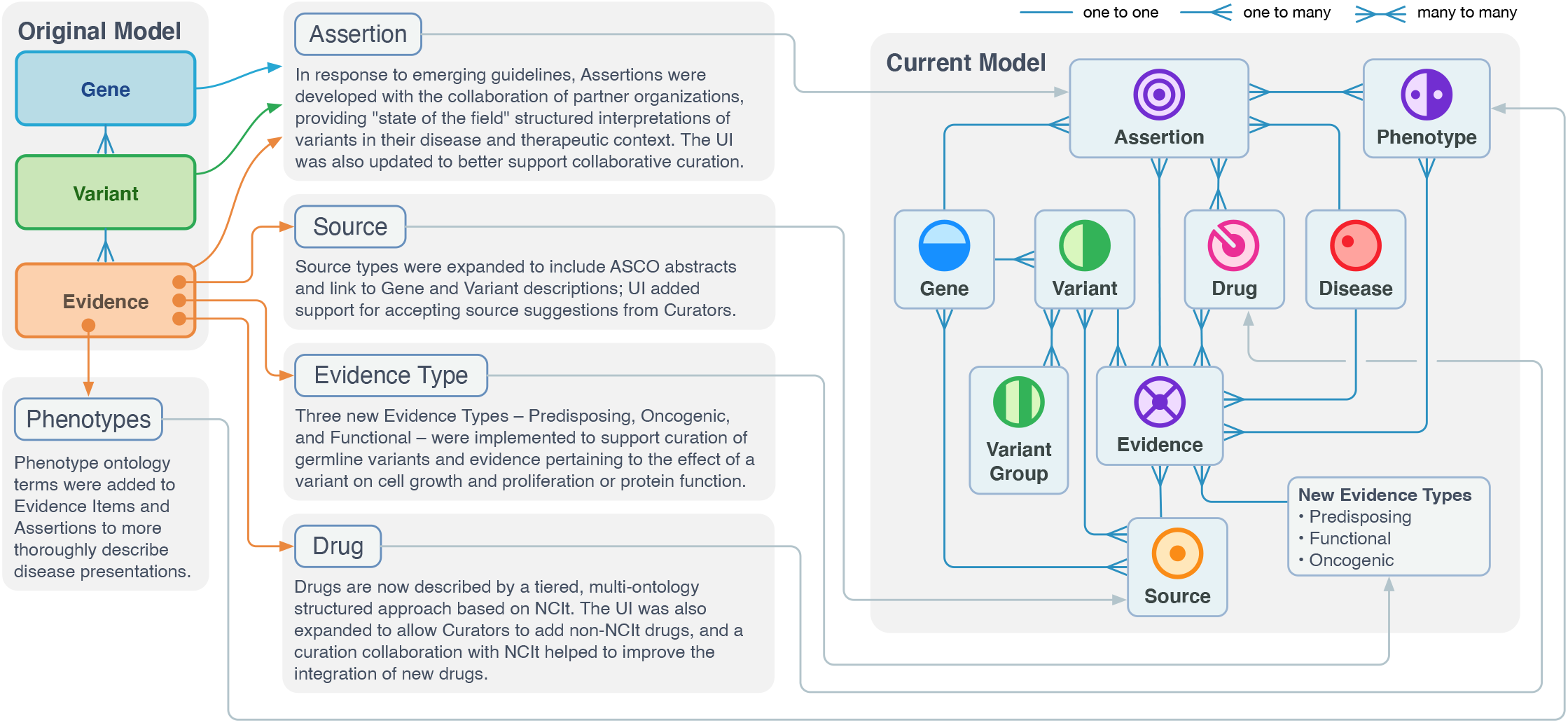
Major feature upgrade to the CIViC Interface. Several upgrades have been made to the CIViC interface since the introduction of the knowledgebase in 2017^1^. Key updates include the introduction of Variant Groups, Assertions, Source Suggestions, and Phenotypes as well as the expansion of Evidence Types and Sources. Many of these features were implemented based on existing collaborations with CIViC users or to align CIViC curation with recognized guidelines. Abbreviations: UI, User Interface.

In 2018, the FDA announced a mechanism for recognition of public human genetic variant databases^18^. A key criterion in the FDA recognition of genetic databases is transparency and public accountability, including description of expert panels and their members^19^. To align with these guidelines, we have implemented Editor COI statements, a formal SOP^2^, and the Organizations feature in CIViC. Curators are associated with specific Organizations and Sub-Organizations to track and display curation activity of various groups (**Figure 5**). Each action performed by a Curator is tagged with their Organization, or, if the Curator is associated with multiple Organizations (or Sub-Organizations), Curators can toggle between their Organizations for any individual action. Currently, CIViC features eleven parent Organizations, the largest of which is ClinGen with 93 members and 5 Sub-Organizations (https://civicdb.org/organizations/2/summary). Organizations have individual pages that display membership, summary statistics, and an activity feed which transparently display contributions and affiliations.

**Figure 5.**
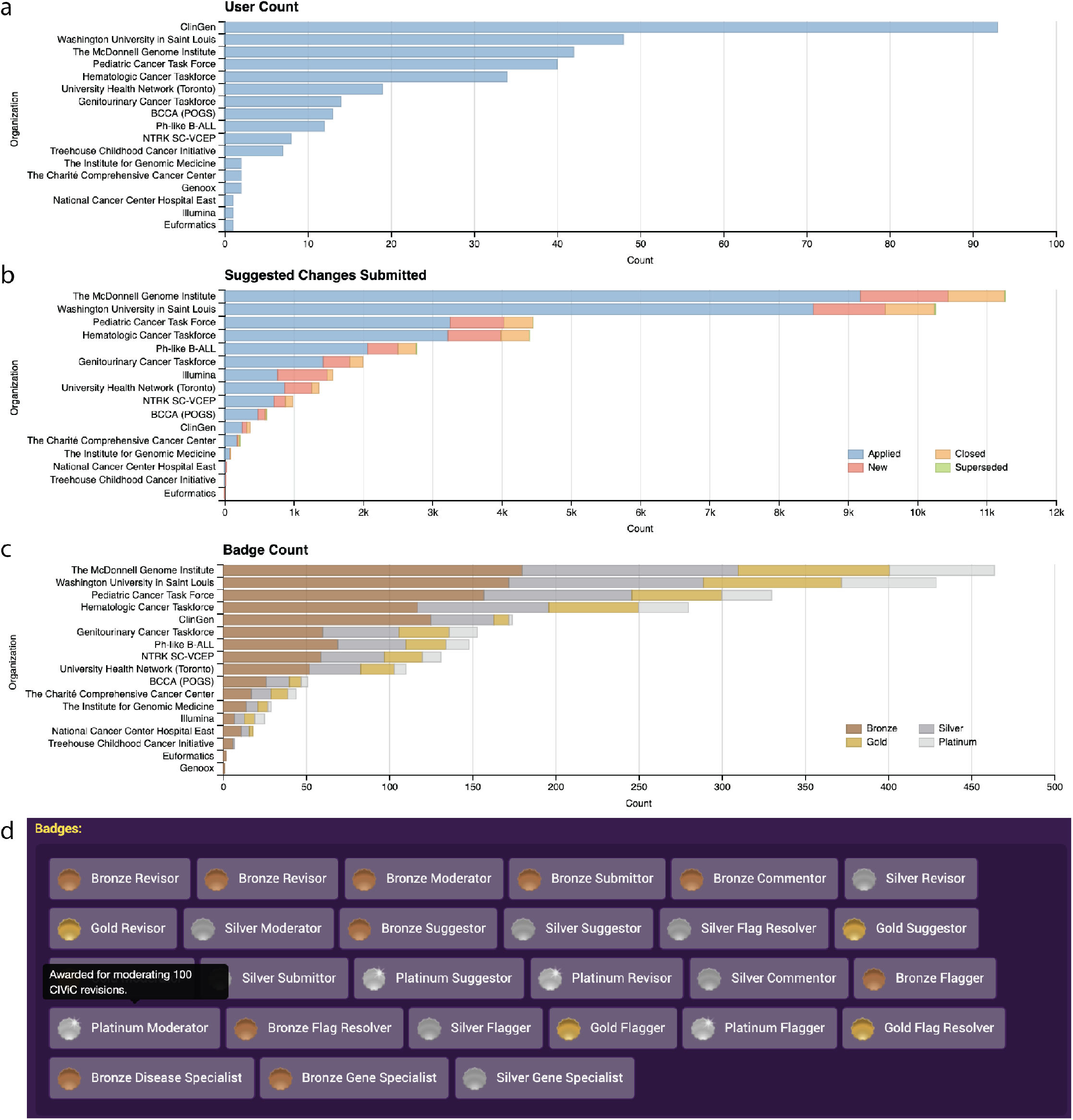
Curator incentivisation through Organizations and Badges. Activities attributed to CIViC Organizations including **a)** Users associated with each Organization, **b)** Changes suggested by each Organization, and **c)** Badge counts for users associated with each Organization. **d)** Example of Badges awarded to Curators as displayed on the User Profile. Badges are assigned from Bronze to Platinum by total actions related to curation and moderation activities within CIViC. Detailed description of the Platinum Moderator Badge is shown in the hover text. Full descriptions can be seen in the user interface for all earned Badges. More statistics can be found at https://civicdb.org/statistics/organizations.

### Collaboration-driven evolution of the CIViC data schema to support new Evidence Types

The evolution of the CIViC data scheme has been as community-driven as the curation itself. Developments have ranged from major overhauls to support emergent guidelines, to adding small use-case-specific features to support external collaborations (**Figure 4**). To obtain feedback and implement changes, curators and developers are routinely engaged through biennial, in-person Hackathon and Curation Jamborees, briefly outlined in the **Supplementary Information**. Specific examples of community-driven features are described below.

At our first Curation Jamboree, the importance of supporting inherited cancer predisposing variants, within the same interface as the subsequent (second hit) somatic variants that lead to cancer development, was discussed in relation to von Hippel-Lindau (VHL) disease. Somatic, inactivating *VHL* variants are the most frequent genetic aberration in clear cell renal cell carcinoma (ccRCC)^20,21^. Meanwhile, Von Hippel-Lindau (VHL) disease is a rare syndrome caused by pathogenic germline variants in the *VHL* gene and is associated with increased risk for ccRCC. Approximately 70% of patients with VHL will develop ccRCC, which remains the leading cause of disease-related mortality^21^. From connections made at this event, a collaboration was undertaken with VHL researchers and clinicians. This led to development of a new Evidence Type – *Predisposing Evidence* – to support detailed curation of germline variants in the CIViC interface. By supporting both germline and somatic variants, CIViC is uniquely situated to propel understanding of the complex interplay between inherited and acquired genetic events in cancer, an area increasingly recognized in clinical guidelines internationally^6,22–24^. To support germline variant evidence and interpretation guidelines^25^, standard procedures were developed for the addition of Human Phenotype Ontology (HPO)^26^ terms and germline interpretation guidelines (in the form of the American College of Medical Genetics/AMP evidence criteria outlined in Richards et al., 2015)^25^ to Evidence Items (**Figure 4**). HPO terms permit the tagging and searching of the underlying phenotypes associated with variants described in the literature. Aggregation of germline evidence is now supported in the form of Assertions that are given ACMG/AMP classifications (e.g., Pathogenic, Likely Pathogenic) where relevant ACMG/AMP evidence criteria (e.g., PVS1, PP1, BS1) are aggregated. As a result of community requests at our Curation Jamboree, and the fruitful collaboration that followed, CIViC now contains the largest known database of VHL disease-associated variants (**Figure 6**, **Supplemental Figure 3**).

**Figure 6.**
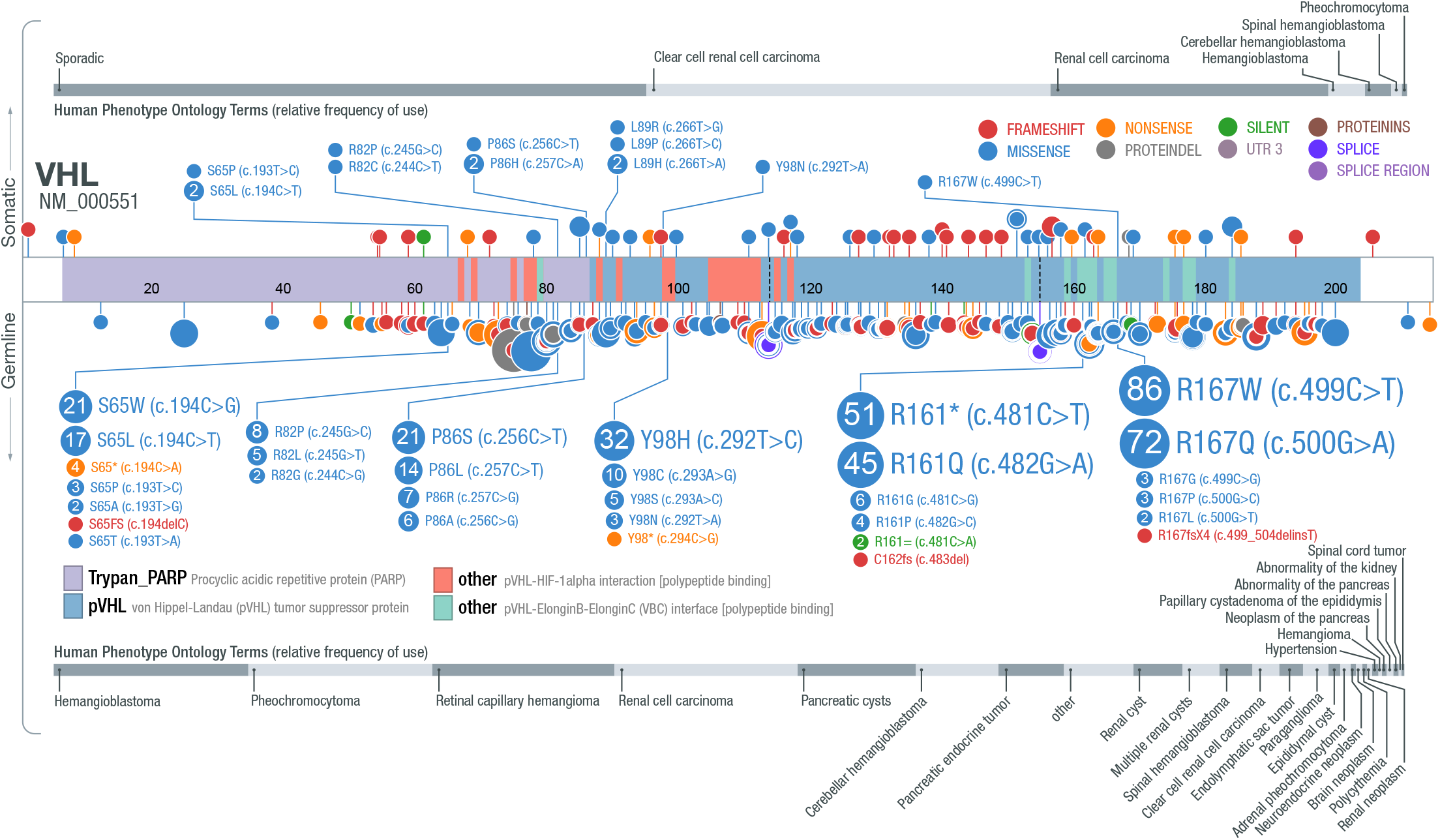
Germline and somatic VHL variants with Evidence Items and associated Human Phenotype Ontology terms. CIViC Evidence Items include VHL variants identified in either somatic (top) or germline (bottom) contexts and displayed using St Jude’s ProteinPaint. Numbers represent the total Evidence Item counts (submitted or accepted) associated with each Variant. Each Evidence Item can have more than one HPO term associated, the distribution of these terms is displayed in grey with bars representing their relative frequency. Only small Variants (single nucleotide variants and small insertion / deletions) with curated coordinates are shown.

Improvement of CIViC structured data has also come from our Hackathon and Curation Jamboree events. Identifying an appropriate ontology to support the Drug field used for Predictive Evidence Items had been a major challenge. At the inception of CIViC, no single ontology encompassed the breadth of drugs and treatments entered into CIViC (from preclinical investigational compounds to FDA-approved therapies), while reducing redundancies by supporting sufficiently curated names and aliases. The Hackathon working group proposed and began implementation of a tiered approach using the NCIthesaurus (NCIt)^28^ as the main source for drug concepts. We normalized 79% of existing Drugs in CIViC in our initial attempt and now use this ontology to automatically search for and normalize new content. To address terms not currently represented in NCIt, we allow non-NCIt Drugs to be entered, and through a more direct collaboration with NCIt, we curate and submit these terms to NCIt for integration on an ongoing basis, enriching both resources (**Supplemental Figure 4**). Consistent feedback and submissions to resources we use, such as the NCIt^28^, Disease Ontology^29^ and ClinGen Allele Registry^30^, promotes a collaborative ecosystem and provides a direct conduit for expert feedback from the needs of the CIViC community to these resources.

Through collaborations with the Variant Interpretation for Cancer Consortium (VICC; cancervariants.org) and ClinGen SC-CDWG, we identified the need to curate evidence pertaining to a variant’s impact on protein function or cellular properties. Large-scale genomic assays, designed to describe the function of numerous variants, allow for the evaluation of rare variants through comparison to established hotspot or targetable counterparts in the same gene^31^. CIViC has set out to more clearly categorize variants based on their ability to induce measurable protein and cellular changes, by modifying the recently described *Functional* Evidence Type^2^ to accommodate the creation of the *Oncogenic* Evidence Type. Both are described in more detail in the following text with additional examples available in the **Supplementary Information**.

Functional Evidence strictly represents the variant’s impact on protein function independent of disease context. Functional Evidence was designed to support fundamental genetic concepts introduced by Mueller’s Morphs^32^: Gain of Function (hypermorphic), Loss of Function (amorphic), Unaltered Function (isomorphic), Dominant Negative (antimorphic), Neomorphic, and Unknown (**Supplemental Figure 5**, **Supplemental Figure 6**). Full inclusion of Mueller’s Morphs allows much more granular representations of protein level effects than those offered by most other resources, which are often limited to gain and loss of function. Functional genomics studies have been specifically designed to query variants for neomorphic and dominant negative properties, which can drive different phenotypic effects and alter recommended treatment courses^33–35^. Our Functional Evidence structure provides the capacity for a thorough categorization of functional genomic results.

The Oncogenic Evidence Type was created to address variant interpretations related to the Hallmarks of Cancer^36^. More specifically, Oncogenic Evidence describes a variant’s role in influencing cancer development through cell growth, clonogenicity, epigenetics, etc. Oncogenic properties are often cell-type dependent, so we require this Evidence Type to be associated with a Disease^37^. Oncogenic Evidence may be used to demonstrate that a variant has properties similar to another variant in the same gene approved for treatment to allow inferences to be made for drug treatment, as outlined in the ESMO Scale for Clinical Actionability of molecular Targets (ESCAT)^38^. Support for Oncogenic evidence is particularly important for rare variants in genes with well-established oncogenic roles (**Supplemental Figure 7**). For example, common oncogenic KRAS variants are found at codons 12, 13, 61, and 146, but rare variants also occur. In these cases, information regarding oncogenicity of these rare variants is important because in colorectal cancer activating KRAS variants are associated with resistance to EGFR inhibitors. CIViC contains Oncogenic Evidence Items (EID9330 and EID9331, **Supplemental Figure 7b**), which describe rare activating in-frame insertions found in colorectal cancer patients, near the common G12 driver variant site^39^. *In vitro* and *in vivo* studies described in these Evidence Items establish the oncogenic role of these rare variants and could be used to influence clinical decision making.

Community engagement and feedback has emphasized the need for supporting curation of abstracts from national meetings where clinical trial results are presented. These often represent the most current results available and may include interim or final clinical trial results that will go unpublished. An evaluation of clinical trial results for breast, lung, colorectal, ovarian, and prostate cancers reported in abstracts from annual ASCO meetings (years 2009-2011) showed that 39% of findings remain unpublished 4-6 years later^40^. Failed clinical trials often provide pertinent information for variant interpretation, but are less likely to be ultimately published. In other cases, regulatory approvals may be based, in part, on evidence only available in conference proceedings. To address the need for curated information derived from ASCO meetings, CIViC has augmented the accepted Source Types to support ASCO abstracts (**Supplemental Figure 8**). Curation procedures recognize this information should be used with caution given the limited access to detailed methodology, and that curation should only reflect the available data. Unfortunately, the licensing and computational accessibility of content from additional peer-reviewed national meeting abstracts remains challenging for broader implementation, though we continue to pursue integration of other Source Types wherever possible, on a case-by-case basis.

### Extension of CIViC software to highlight and integrate other variant resources

In addition to curation-driven collaborations, we continue to expand our software development collaborations. Manually providing depth and breadth of coverage of the ever-expanding biomedical literature is challenging for highly specialized curation tasks, such as identifying relevant cancer variants. Comparisons of cancer variant knowledgebases, through collaboration between the VICC^7^ and CIViC Editors^41^, have demonstrated a surprising dearth in publication overlap. To address this gap, colleagues at Canada’s Michael Smith Genome Sciences Centre at BC Cancer leveraged experienced CIViC Editors to train a natural language processing model called CIViCmine to identify high-priority publications for CIViC and other cancer variant knowledgebases (http://bionlp.bcgsc.ca/civicmine/)^9^. Ongoing efforts continue to expand the functionality and improve the integration of CIViCmine.

Other critical collaborative projects have been expanded by outreach to other resources. The incorporation of the ClinGen Allele Registry^30^ enables users to connect manually curated (genomic) CIViC Variant Coordinates to their preferred genome build or transcript reference using the Allele Registry link on CIViC Variant pages or by retrieving the Canonical Allele ID through the CIViC API. Users evaluating variants via the ClinGen Allele Registry are similarly offered links back to CIViC. Analogous collaborations with the developers of St. Jude’s ProteinPaint tool^42^ have led to bidirectional links from CIViC to ProteinPaint, providing a visual representation of Variants in CIViC with curated Coordinates. ProteinPaint users are directed in the interface to curated CIViC Evidence for their variants of interest. Additional collaborative products that have come from the CIViC knowledgebase can be found in **Supplemental Table 1**.

As the CIViC data schema and connections have expanded, additional documentation for Curators and software developers was necessary. Migration of our help documentation to a dedicated interface (hosted by readthedocs) lowers the maintenance cost for this documentation, a much-needed improvement that coincided with the development of a formalized standard operating procedure^2^. Additional warnings and default page states have been introduced, which have become mainstays of the CIViC user experience, to emphasize higher-quality content and notify users of pending changes or unmoderated content. We have also provided faster, lower burden curation tasks, such as *Flags* to quickly draw Editor attention to potential errors (**Supplemental Figure 1**), Source Suggestions to recommend content for curation (**Supplemental Figure 8**), improved Advanced Search options (**Supplemental Figure 9**), and a redesign of the Variant interface (**Supplemental Figure 10**, **Supplementary Information**). Bioinformaticians and developers have taken advantage of the ease of CIViC’s API for integrations, with multiple groups integrating CIViC with no direct interaction with our team (**Figure 1**, **Figure 2**). In addition to the API, data releases are now made available through monthly and nightly TSV and VCF files (https://civicdb.org/releases). TSV files are available for each of the major CIViC entities (Evidence Items, Assertions, Variants, and Genes). VCF files summarize CIViC Variants with curated coordinates and include annotations with data of submitted and accepted Evidence Items and Assertions linked to each Variant. These VCF files can be used to annotate patient variant calls with data available from CIViC for rapid clinical variant interpretation. Hackathon events have also led to custom data formats and collaborative development to incorporate CIViC data into external resources such as NDEx^43^, myvariant.info^44^, WikiData^45^, and openCRAVAT^46^. For more programmatic approaches, CIViCpy^10^, a software development kit enabling advanced CIViC queries, was developed for users familiar with Python.

### CIViC as an educational and training resource

An increasing demand on the scientific community is the education and training of the next generation of biocurators, variant analysts, and geneticists^47–49^. By making CIViC available without any installation requirements beyond a web browser, and permitting any registered user to be a CIViC Curator, CIViC’s curation and variant interpretation interface has a low barrier for access, which proves useful in educational settings and workshops. After initial training on one of our staging servers, trainee submissions are evaluated and moderated by Expert Editors, maintaining the quality of the CIViC knowledgebase while providing feedback to trainees. Features such as the Source Suggestions queue (https://civicdb.org/curation/sources) (**Supplemental Figure 8**) provide a pre-selected list of potential PMIDs for curation that can be searched by gene, variant, disease or publication year. At the same time badges incentivize trainees by rewarding curator activity (**Figure 5c,d**). Training in CIViC curation promotes direct engagement with the clinical literature, develops skills in extracting evidence from the published literature, and provides interaction with clinical experts through the interface independent of time zone or physical location. Through summer internship programs, research collaborations, and formal courses, CIViC has been used as part of the training for undergraduate and graduate students with an interest in oncology or research in genetics. CIViC interns and undergraduate trainees have gone on to nursing, medical, and graduate school training programs. An open but rigorous variant interpretation resource such as CIViC can facilitate community engagement and provide mechanisms for the education of contributors, which ultimately produces higher quality contributions while supporting the advancement of the community at large.

## Discussion

The CIViC knowledgebase exemplifies how open-access platforms can permit widespread adoption and dissemination of cancer variant interpretations. The six founding principles of CIViC (interdisciplinary, community consensus, transparency, computationally accessible, freely accessible and open license) have proven to be a robust foundation for rapid adaptation and widespread adoption. Contributions to CIViC can be made by every stakeholder in the field, including scientists, physicians, students, and patients. Engagement from diverse communities improves the ability to curate information quickly and the use of structured data models allow for all users to contribute meaningfully, regardless of their background. Through crowdsourcing and targeted engagement techniques, we have successfully recruited Editors and Curators from around the globe. Editors and Curators continue to be actively recruited to the CIViC knowledgebase. Interested individuals can find more information on joining the community through the CIViC help documents (docs.civicdb.org). As we initially designed and have demonstrated, even minor contributions to the database compound into a global impact that improves the way we access and interpret cancer variant data.

As clinical sequencing of cancers is adopted by increasing numbers of clinics, companies, and hospitals, the need for curated information to guide clinical variant interpretation continues to grow. This is further compounded by the exponential growth in medical literature, which greatly surpasses the ability of any one institution or group to assimilate. This bottleneck emphasizes the need for both open-access resources and community-generated contributions. By creating a resource with a low barrier to entry paired with expert moderation, CIViC promotes community engagement that has the potential to scale with the medical literature. Following the model of widely-adopted bioinformatic tools used for next generation sequencing analysis, promoting CIViC as an open-access and programmatically-accessible resource has made a significant contribution to this effort. For example, CIViC has been widely adopted internationally^50^, and embraced by the patient community (including support from the VHL Alliance) to become the largest, publicly available *VHL* variant knowledgebase. VHL literature and evidence curated in CIViC have also become a valuable resource for the ClinGen VHL germline Variant Curation Expert Panel (VCEP) to develop their classification rules and as an external source of evidence when curating germline variants in the ClinGen Variant Curation Interface (VCI)^27^. CIViC is not designed to replace the critical role of clinical geneticists in variant interpretation, but provides a tool to assist professionals in performing their clinical responsibilities.

The CIViC Ecosystem relies on the considerable volunteer efforts of Curators and Editors, resources it has incorporated, and user feedback. In turn, CIViC contributes to the ongoing development of this community through improved open-access data distribution, creation of educational materials, and providing feedback and curation to external resources. The acknowledgement and support of this community is critical to CIViC’s success. This work remains unfinished. There is a critical need fors more expert Editors, to moderate the contributions of Curators, and keep pace with the ever expanding medical literature. The use of the CIViC platform for curation enables expert working groups from widely disparate time zones to collaborate asynchronously. However, despite engaging a diverse team of Curators, ongoing efforts are needed to engage not only more community Editors, but Editor representation outside of North America and Europe.

Development work continues to address more complex issues, such as: variant relationships and combinations with clinical significance, improved structural representation of complex variants (e.g., copy-number and structural variants), and large-scale variants that impact more than one gene. Such changes require substantial development efforts to adapt the database schema and the user interface. As this resource expands, we are committed to ensuring this work remains free and open to all, without fees or restrictions.

We have shown that augmenting the remarkable efforts of biocurators and geneticists through structured and open data is a viable and robust way forward for cancer variant interpretation, thereby enabling the democratization of genomics in patient care. This massive collaborative effort amplifies the skills of curators, bioinformaticians and developers to produce a knowledgebase ready to adapt to the changing needs of the field of cancer variant interpretation.

## Acknowledgements

We want to thank the CIViC community and specifically acknowledge and thank the following colleagues for their contributions to this work and helpful feedback on the manuscript: Saleh Albanyan, Sydney Anderson, Garrett Bullivant, Laura Corson, Justin Guerra, Chimene Kesserwan, Ian King, Geoff Lyle, Sharon Plon, Nathan Schachter, Dmitriy Sonkin, Kristen Sund, Gregory Stupp, Marta Szybowska, Anna Tanska, Lee Trani, Brian Walsh, and Amber Wollam. We also thank the tireless efforts of our omniscient and beloved civic-bot.

The CIViC project was supported by the NCI under Award Number U01CA209936 and U24CA237719 including funding supplements from the Cancer Moonshot and Childhood Cancer Data Initiative (CCDI). CIViC was also supported by the Washington University Institute of Clinical and Translational Sciences grant UL1TR002345 from the National Center for Advancing Translational Sciences (NCATS) of the National Institutes of Health (NIH). CIViC is also supported by Children’s Discovery Institute (CDI) of the St. Louis Children’s Hospital and Washington University School of Medicine. This research benefited from the use of credits from the National Institutes of Health (NIH) Cloud Credits Model Pilot, a component of the NIH Big Data to Knowledge (BD2K) program. Additional support was received from the Google Big Query and Amazon Web Services Open Data projects. Research into germline VHL has been supported by a VHL Alliance Research Grant. **Andreea Chiorean** was supported by the Starbucks Clinical Genetics/Genomics Research Studentship Award 2018. **Damian T Rieke** is a participant in the Berlin Institute of Health - Charité Clinical Scientist Program funded by the Charité -Universitätsmedizin Berlin and the Berlin Institute of Health. **Deborah I. Ritter**, **Aleksandar Milosavljevic**, and **Ronak Y. Patel** were supported by the Clinical Genome Resource (ClinGen) grant U41HG009649 from the NHGRI. **Raymond H Kim** was supported by the Bhalwani Family Charitable Foundation. **Steven JM Jones** is UBC Canada Research Chair in Computational Genomics. **Alex Handler Wagner** was supported by the NHGRI under award number K99HG010157. **Obi Lee Griffith** was supported by the National Cancer Institute of the NIH under Award Number K22CA188163. **Malachi Griffith** was supported by the National Human Genome Research Institute (NHGRI) of the National Institutes of Health (NIH) under Award Number R00HG007940.

