## Supplemental for "Evolution of the open-access CIViC knowledgebase is driven by the needs of the cancer variant interpretation community"

**Supplementary Information** **1**
CIViC Hackathon and Curation Jamborees 1
Functional and Oncogenic Evidence Examples 2
Relevant links to resources mentioned in the main text 2
**Supplementary Figures** **4**
Supplemental Figure 1. Range of curation activities within CIViC 4
Supplemental Figure 2. Growth of Curator base Acquisition of new curators 5
Supplemental Figure 3. CIViC Evidence Grid displays somatic and germline evidence for VHL S65L 6
Supplemental Figure 4. New ‘Add Drug’ Interface 7
Supplemental Figure 5. Mapping of CIViC Functional Evidence to the classical genetic concepts of Mueller’s Morphs 8
Supplemental Figure 6. Functional Evidence Item examples 9
Supplemental Figure 7. Oncogenic Evidence Item examples 10
Supplemental Figure 8. Integration of Sources to CIViC over existence of the knowledgebase 11
Supplemental Figure 9. Enhancement of the Advanced Search Page 12
Supplemental Figure 10. Variant interface, then and now 13
**Supplementary Tables** **14**
Supplemental Table 1. Peer-reviewed publications associated with CIViC collaborations 14
**References** **15**

**Supplementary Information**

*CIViC Hackathon and Curation Jamborees*

The CIViC team has organized and hosted biennial "Hackathon and Curation Jamboree" events, attended by multiple stakeholders in the cancer variant interpretation community (civic.readthedocs.io/en/latest/about/meetings.html). These events are open to the community and occur in the style of an “unconference” where topics are driven by attendees[1]. Attendees freely join breakout sessions and pursue topics of their choice while bringing back results or conclusions to the group as a whole at the end of the day or as part of the final session. To maximize the involvement in groups and presentations as well as interactions amongst participants, the attendee list is kept relatively small (registration capped at <50) and involves a few scheduled social events and meals over the 2-3 day period.

The first event was held at the Netherlands Cancer Institute (NKI) Nov 30 - Dec 2, 2016 as part of the “NGS in Molecular Pathology Symposium”. The second was held at Scripps Research Institute, La Jolla, CA Oct 15 - Oct 16 2018, as an adjunct meeting ahead of the American Society of Human Genetics (ASHG) annual meeting. A third was planned for Fall of 2020 but postponed. These meetings focused on topics in the CIViC open-source code base, curation in CIViC, and issues of curation for the cancer variant community in general. Specific topics and breakout sessions from these meetings can be viewed on GitHub (github.com/griffithlab/civic-meeting). For example, topics in 2016 included implementing badges and curation incentives into CIViC, incorporation of pharmacogenomic and predisposing biomarkers in CIViC, and capturing the complexities of therapeutic responses in preclinical models in a variant knowledgebase. Topics in 2018 included exporting CIViC data to variant call format (VCF) files, making CIViC networks available in NDEx, systematic evaluation of somatic variant oncogenicity, and best practices in annotating copy number variants (CNVs) and fusions in knowledgebases. These sessions have brought together individuals with a wide range of experience levels to produce collaborative efforts such as resource integrations (e.g., NDEx, CRAVAT, NCI thesaurus), data modeling and exporting, collaborative curation efforts (e.g., VHL as described in the text) and involvement in external organizations (e.g., Somatic Cancer Clinical Domain Working Group [https://clinicalgenome.org/curation-activities/somatic/][2], Variant Interpretation for Cancer Consortium’s (VICC) Knowledge Curation and Interpretations Standards working group [cancervariants.org/wg/kcis/]).

*Functional and Oncogenic Evidence Examples*

Functional Evidence Items in CIViC are designed to capture evidence describing altered protein function due to the presence of a variant. This evidence type does not capture variant effects on cellular behavior, or assess the status of the variant as an oncogenic driver, and does not take disease into account. Functional Evidence will generally describe the results of *in vitro* studies (**Supplemental Figure 6**).

Oncogenic Evidence Items primarily capture *in vitro* evidence, but this Evidence Type focuses on the downstream oncogenic effects of the alteration, regardless of the type of underlying functional change (e.g., loss of function, dominant negative) that led to those oncogenic effects. **Supplemental Figure 7**)

CIViC has a mechanism to capture the observation of a variant in a somatic tumor for assessment as a potential driver. Somatic tumors often harbor many variants — ranging from oncogenic to passenger variants, with many variants having uncertain significance. However, not all variants observed in somatic tumors meet the clinical threshold for inclusion in CIViC. To avoid the accumulation of variants with no established role in cancer, simple observations of a variant in a cancer are not entered into CIViC without substantial additional evidence or support. Therefore, Oncogenic Evidence documented in CIViC should meet stricter criteria for clinical importance, which are outlined in our help documentation (docs.civicdb.org).

In cases where a patient harbors a somatic variant in a gene strongly associated with a germline condition (e.g., *VHL* with Von Hippel Landau [VHL] disease, or *MSH6* with Lynch Syndrome), a Case Study (Level C) Oncogenic Evidence Item may be entered into CIViC, with a one star Evidence Rating. For example, **Supplemental Figure 7c** depicts an Evidence Item for a somatic *VHL* variant seen in a case of sporadic renal cell carcinoma. If the variant was seen in a germline context, it would be entered as predisposing for VHL disease. However, in all instances of Oncogenic Case Study Evidence, the Disease field contains the disease observed in the patient (in this case renal cell carcinoma) rather than the syndrome or disorder associated with the variant in a germline context.

*Relevant links to resources mentioned in the main text*

CIViC interface
- civicdb.org

CIViC help documentation
- docs.civicdb.org

ClinGen Somatic Cancer Clinical Domain Working Group (SC-CDWG)
- https://clinicalgenome.org/curation-activities/somatic/

Variant Interpretation for Cancer Consortium
- https://cancervariants.org/

CIViCmine
- http://bionlp.bcgsc.ca/civicmine/

Example CIViC Organization with Sub-Organizations (ClinGen)
- https://civicdb.org/organizations/2/summary

CIViC API documentation
- https://docs.civicdb.org/en/latest/api.html

CIViC data releases (VCF, TSV
- https://civicdb.org/releases

CIViC Source Suggestions queue
- https://civicdb.org/curation/sources

**Supplementary Figures**

**Supplemental Figure 1. Range of curation activities within CIViC**

There are a variety of actions that can be performed within the CIViC interface that range in time burden for Curators. Comments and flags are the least burdensome activities whereas the generation of new Evidence Items and Assertions represent high time and knowledge requirements. Other activities, including suggesting sources or revisions, can be performed with moderate burden on Curators.

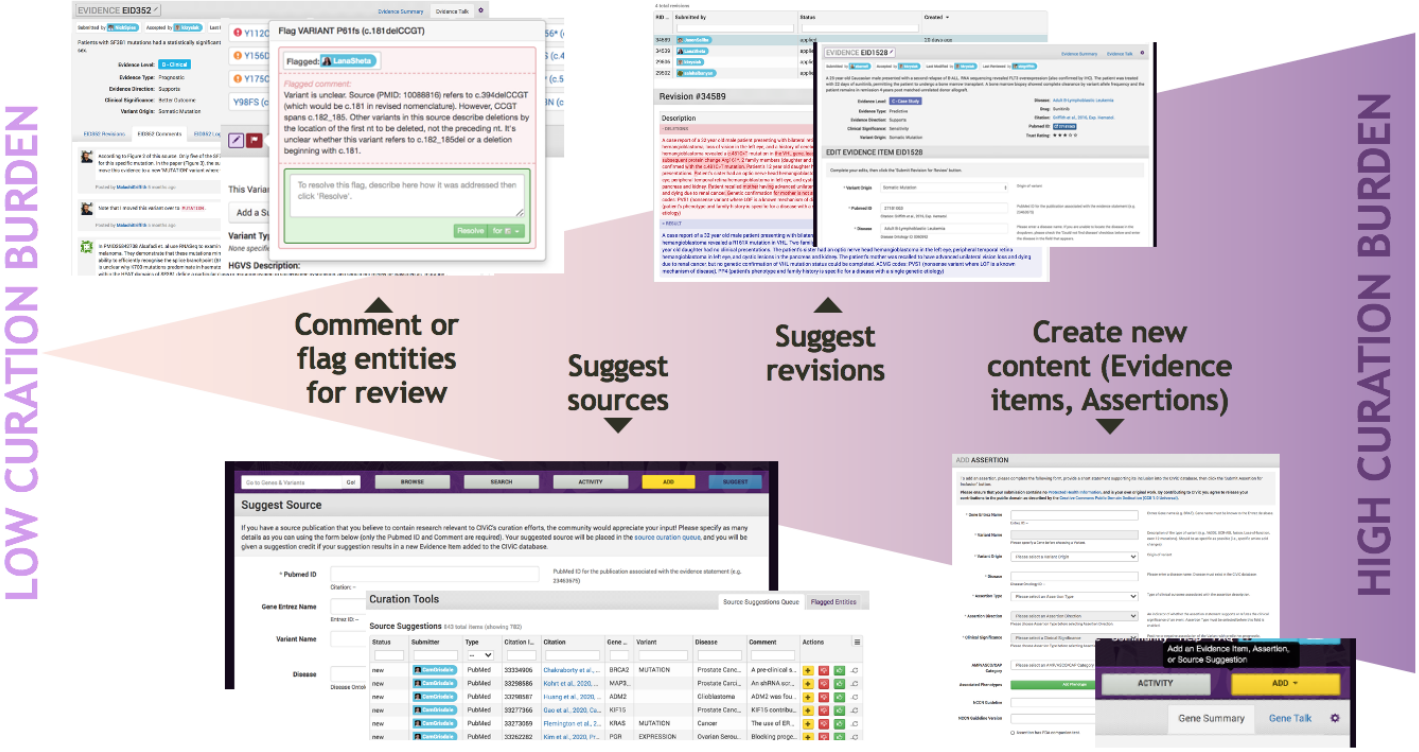

**Supplemental Figure 2. Growth of Curator base Acquisition of new curators**

The growth of CIViC's Curator base since its initial launch in 2015. Curators are defined as users who created an account and performed Curator actions in the interface such as adding Evidence Items, suggesting changes, commenting, or adding Source Suggestions. Total contributing curators are shown in purple. White dots represent the number of new curators added in each calendar month.

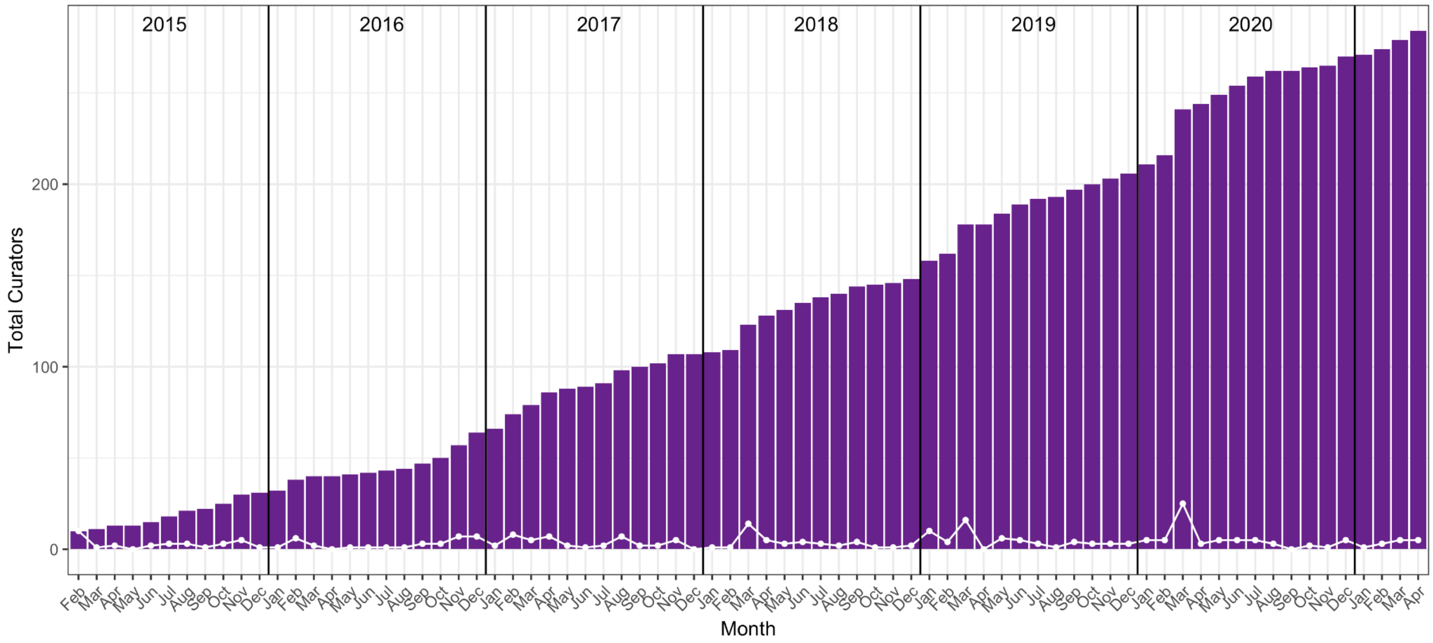

**Supplemental Figure 3. CIViC Evidence Grid displays somatic and germline evidence for VHL S65L**

Screenshot of the CIViC user interface showing the VHL Variant S65L being of rare germline (EID5159), unknown (EID6644) and somatic (EID6131, EID6890) origin in Evidence Items alongside each other in the Evidence Grid. EID6644 is a Level C - Case Study with associated description and HPO terms (https://civicdb.org/links/evidence/6644). The status of this variant is presumed to be germline due to clinical criteria for Von Hippel-Lindau disease, but unconfirmed; therefore the variant origin of this Evidence is Unknown. *EID = Evidence Item ID*

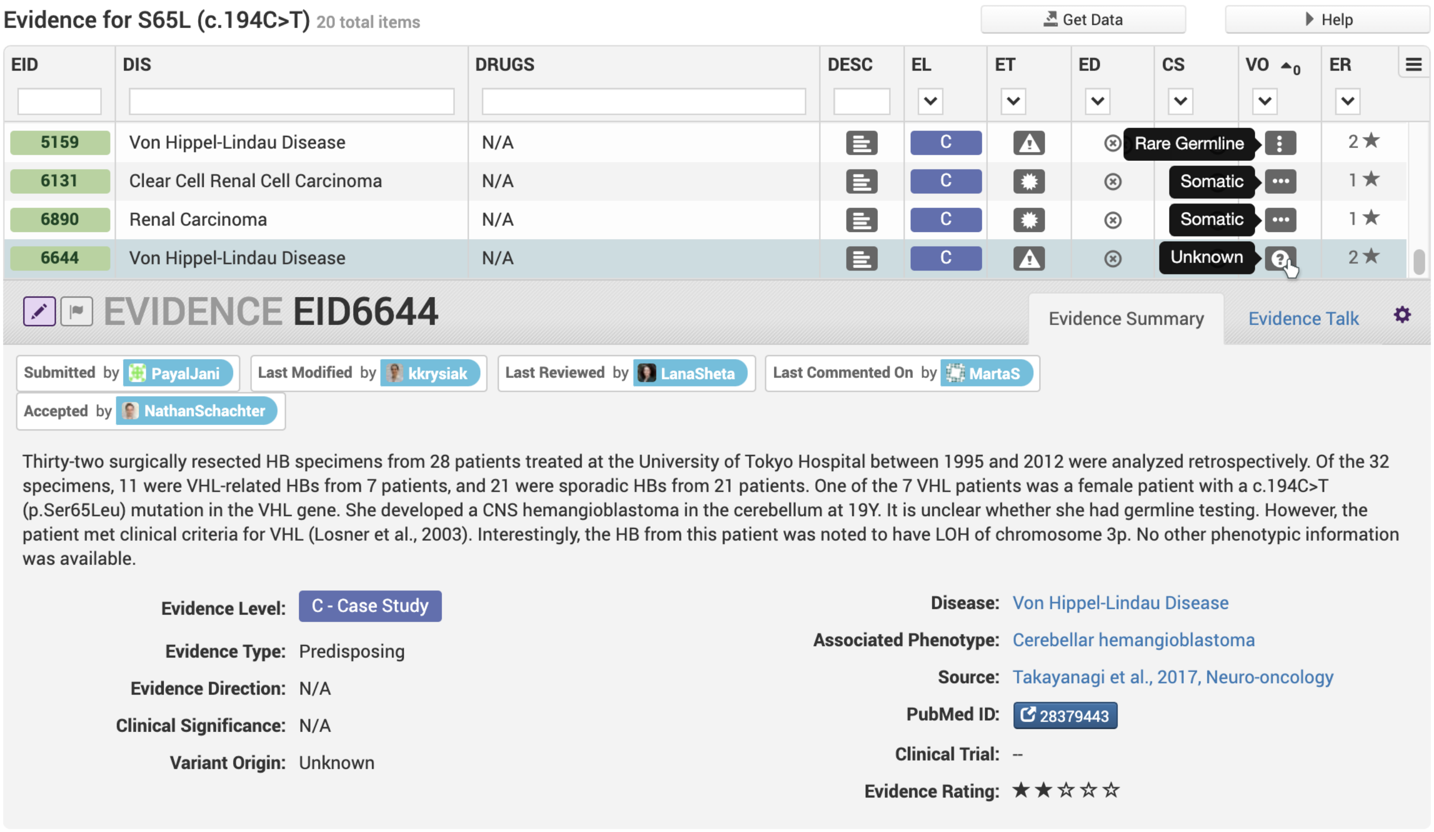

**Supplemental Figure 4. New 'Add Drug' Interface**

The NCI Thesaurus (NCIt) ontology is used to normalize drugs and drug aliases in CIViC. **a)** All Drugs in CIViC and the imported NCIt subset* are searchable when entering or editing Drug Names in CIViC. Evidence items may also be associated with more than one Drug that is used as Substitute, Combination or Sequential treatments in a trial or experiment. **b)** Not all experimental compounds and Drug Aliases needed by CIViC Curators are in NCIt. In order to capture these terms, users are still able to add drug concepts not currently in NCIt via the "Add Drugs" form. **c)** If the user mistakenly tries to create a drug that already exists in CIViC, a warning is supplied and the appropriate Drug can be applied to the Evidence Item. **d)** Display for successful addition of a Drug to CIViC. This new Drug will be subsequently submitted to NCIt but CIViC Curators to improve its use in both resources.

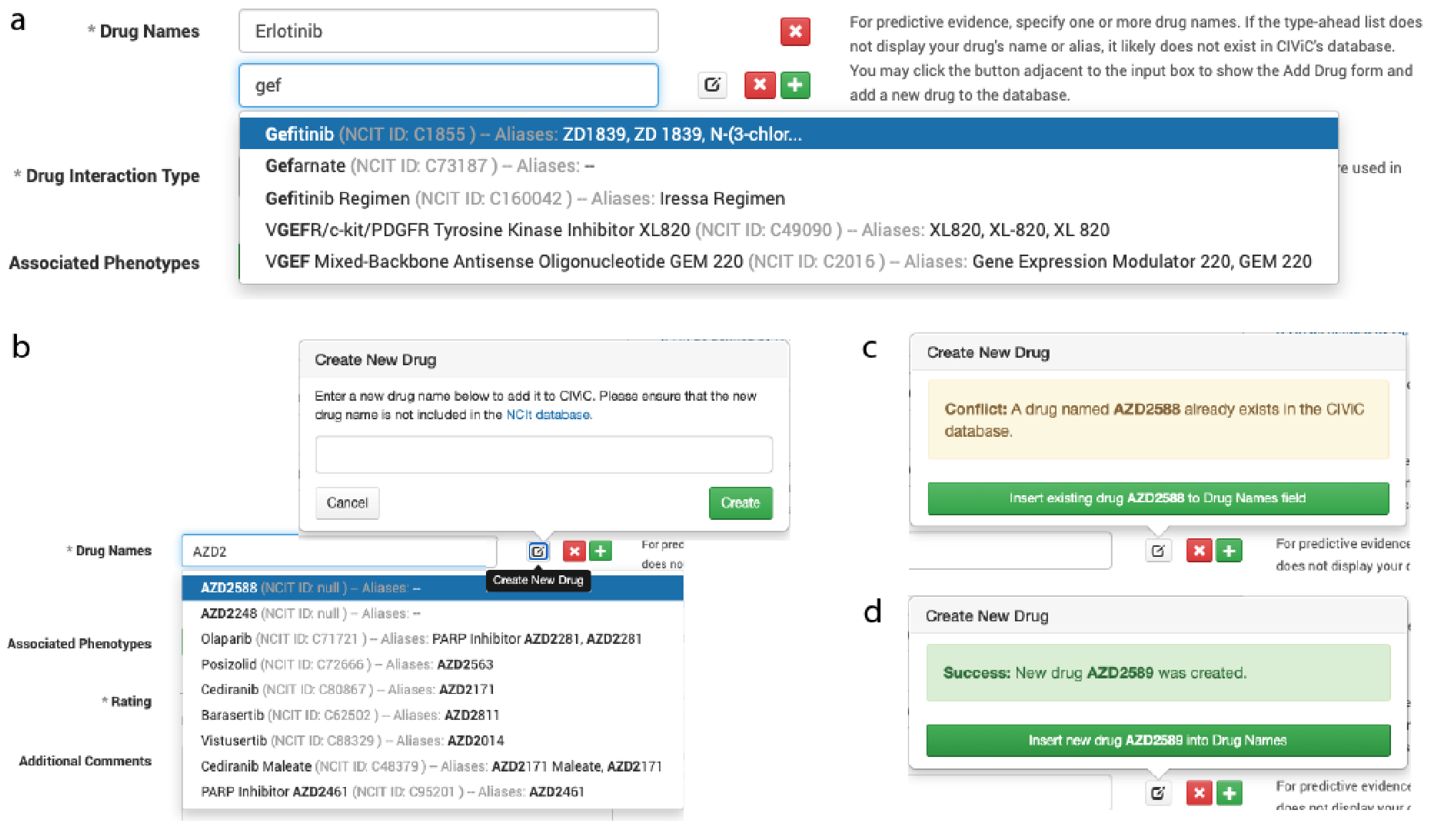


**CIViC imports the following NCIt semantic types: 'Pharmacologic Substance', 'Pharmacological Substance', 'Clinical Drug', 'Therapeutic or Preventive Procedure', 'Hazardous or Poisonous Substance'*

**Supplemental Figure 5. Mapping of CIViC Functional Evidence to the classical genetic concepts of Mueller's Morphs**

CIViC Functional Clinical Significance annotations are derived from the classical Mueller Morphs[3], represented here on a relative variant effect scale to illustrate the spectrum of clinically-relevant alterations a variant can have on protein function. These functional nuances are captured as Evidence and combined with other Evidence Types to support Assertions - variant interpretations with Predictive, Diagnostic, Prognostic, or (cancer) Predisposing associations.

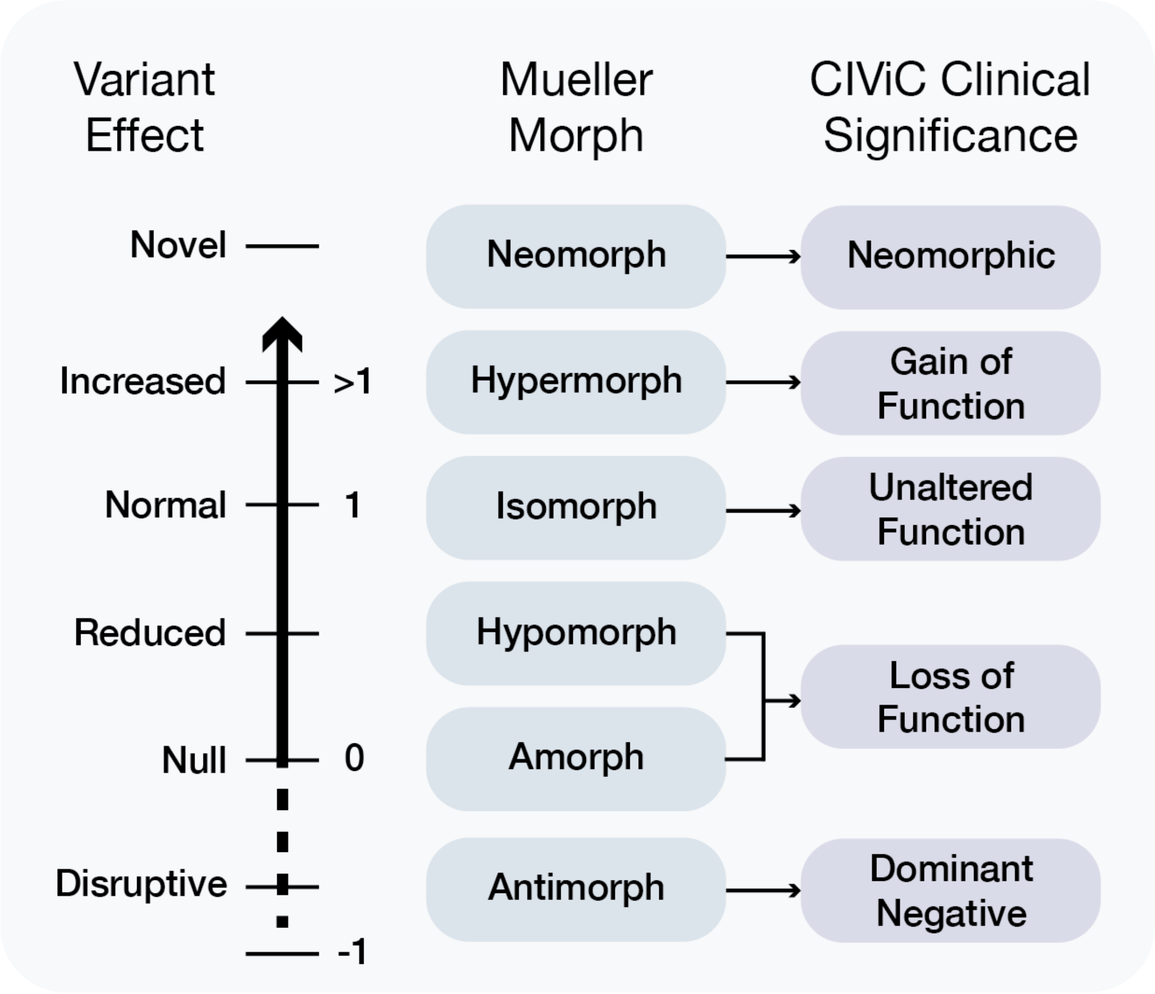

**Supplemental Figure 6. Functional Evidence Item examples**

Examples of curated, moderated and subsequently accepted Functional Evidence Items (EIDs) taken from the CIViC knowledgebase, as displayed in the user interface. **a)** Functional Evidence Item describing preclinical work demonstrating a gain of function effect associated with the non-canonical BRAF A728V variant. **b)** Functional Evidence Item describing a protein activity assay indicating temperature-sensitive loss of function associated with TP53 A161T. **c)** Functional Evidence Item describing preclinical assays which indicate a dominant negative effect associated with the TP53 R248Q variant.

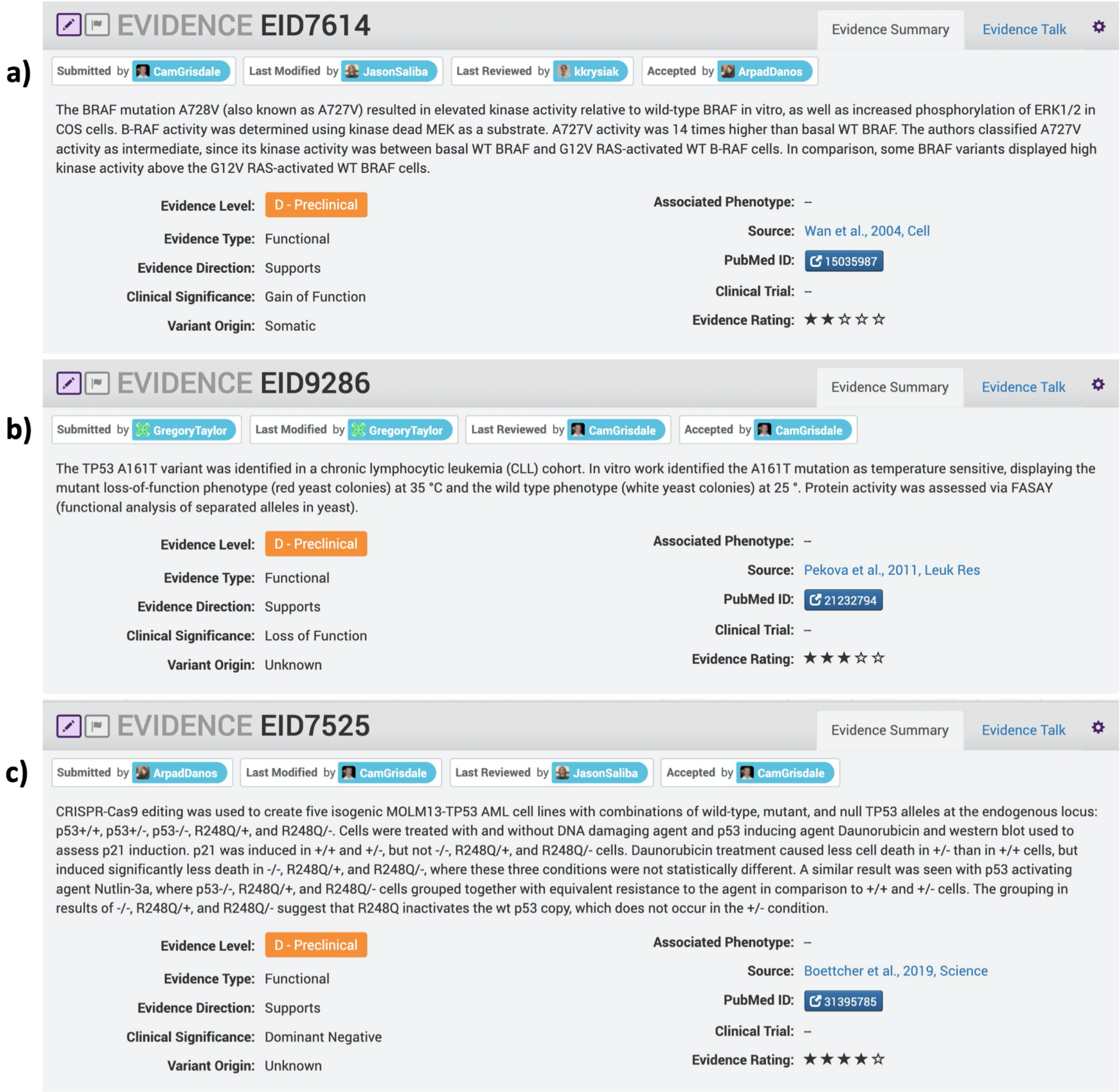

**Supplemental Figure 7. Oncogenic Evidence Item examples**

Examples of curated, moderated and subsequently accepted Oncogenic Evidence Items (EIDs) taken from the CIViC knowledgebase, as displayed on the user interface. **a)** Oncogenic Evidence Item describing preclinical work characterizing density-independent cell growth (foci formation) associated with KRAS Q61H transfection. **b)** Oncogenic Evidence Item describing experiments in cell lines and mice on a rare KRAS in-frame insertion variant discovered in a colorectal cancer patient cohort. The rare KRAS variant is able to promote colony formation in culture and tumor growth in nude mice comparable to a positive control KRAS G12V variant. When compared to controls (G12V and wildtype), these experiments support an oncogenic effect caused by this rare KRAS G10_A11insG variant. **c)** Oncogenic Evidence Item describing a case study with a sporadic renal cell carcinoma patient harboring a somatic *VHL* frameshift variant c.163_164delGA.

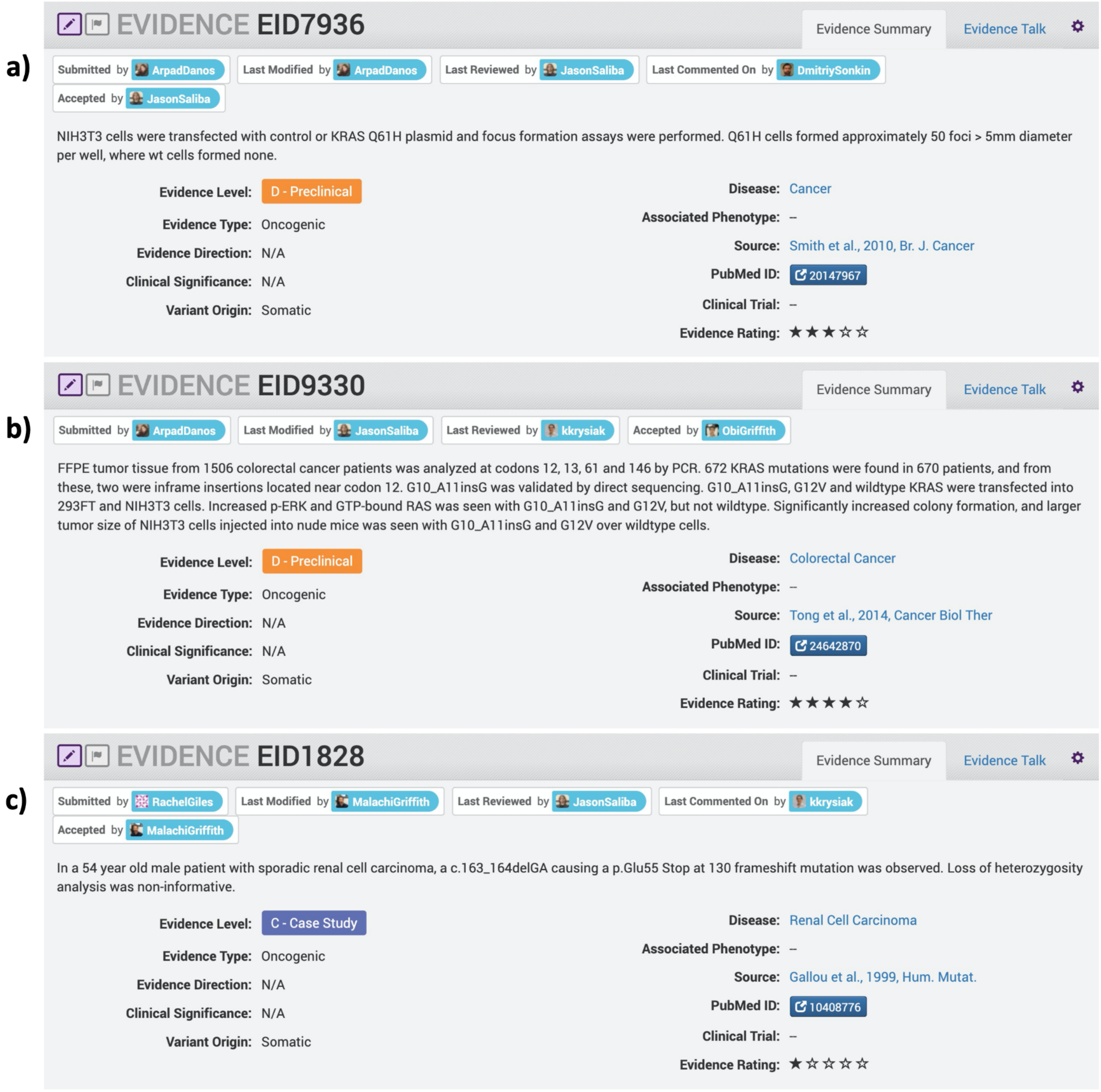

**Supplemental Figure 8. Integration of Sources to CIViC over existence of the knowledgebase**

**a)** This density plot shows monthly counts of unique Sources added, each associated with newly curated content, since the launch of the web interface. A single Source in CIViC may be associated with multiple Evidence Items (e.g., evidence for different Evidence Types or Variants). Each publication is only counted when it is first added to the CIViC knowledgebase. Peer-reviewed, PubMed-indexed literature (purple) was the first Source Type supported in CIViC. The ability to use American Society of Clinical Oncology meeting abstracts as a Source Type was launched in mid-2019 (green). Black dots indicate the accumulation of Sources in the Source Suggestion Queue. **b)** Screenshot of the Source Suggestions Queue (https://civicdb.org/curation/sources) where Curators can quickly recommend publications for curation. To support low-effort contributions, only the publication type (PubMed or ASCO abstract), Citation ID and a comment are required for entry. Hover text shows a comment provided by the Curator at submission. The displayed grid is filtered to citations from the year 2021. Additional sorting and filtering can be performed on any of the displayed entities to enable quick searching for diseases, genes, etc of interest.

a

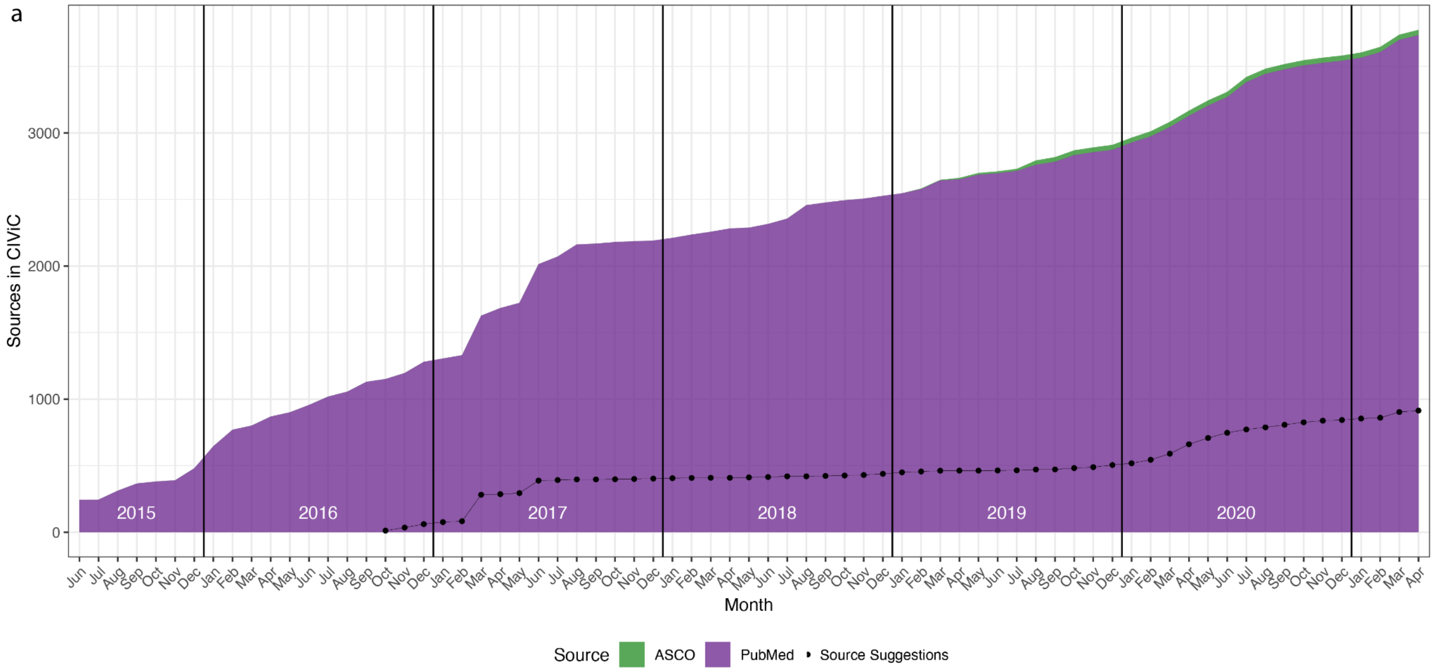


b

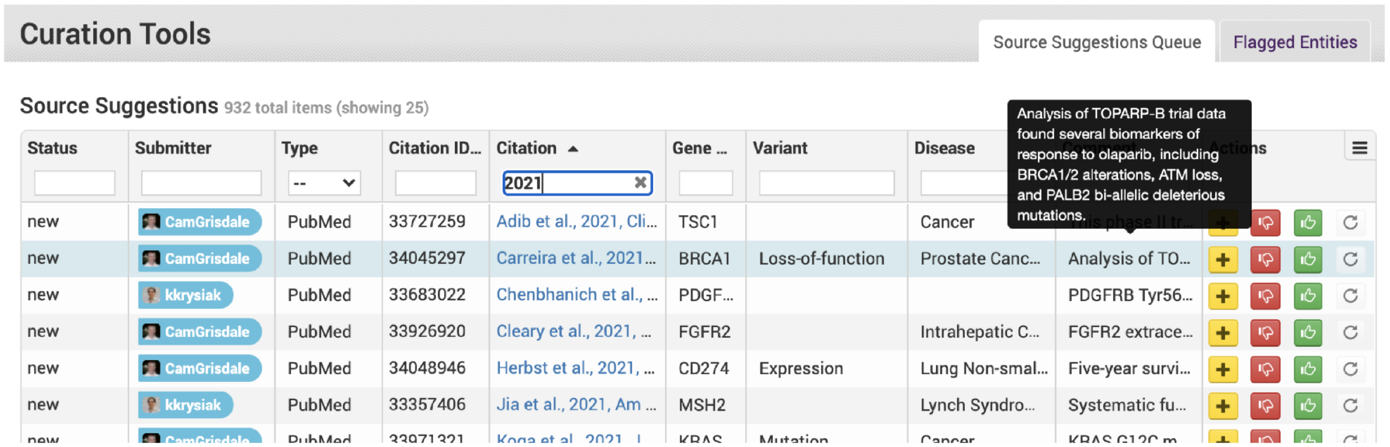

**Supplemental Figure 9. Enhancement of the Advanced Search Page**

Portions of the original Advanced Search Page circa 2016 are shown (top) for comparison to the current Advanced Search Page. **a)** Expanded search options now include new entities (i.e., Assertions) and Suggested Changes. **b)** We have added the ability to search for additional fields, including Human Phenotype Ontology (HPO) terms, Source type (ASCO, PubMed), and Submitter Organization, and create complex queries. **c)** Integration of the NCI thesaurus as a drug ontology allows searching by alias while normalizing Drug names. **d)** Drug combinations are now supported, allowing more than one drug to be associated with an Evidence Item and their relationship labeled as Combinations, Substitutes, or Sequential.

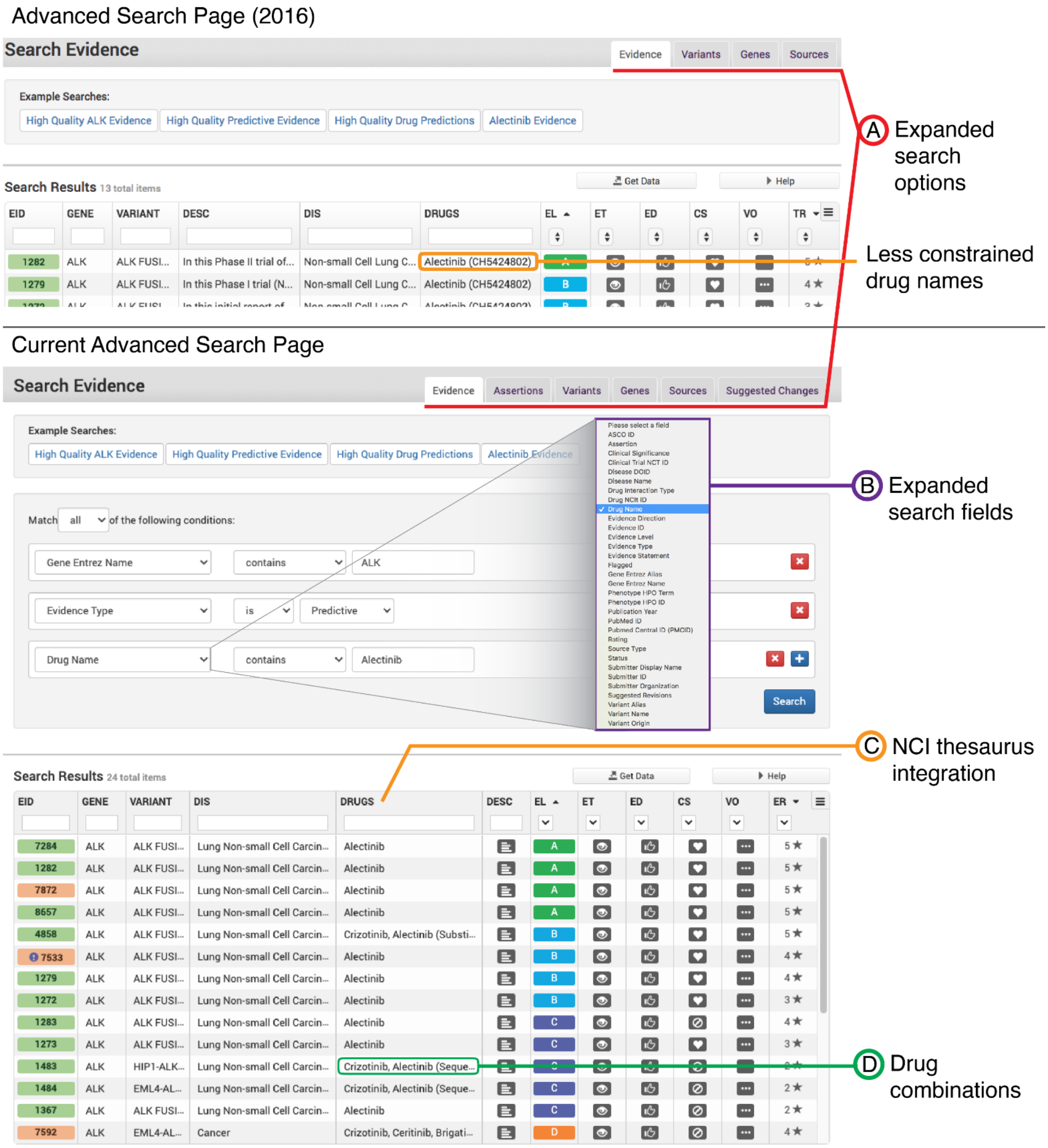

**Supplemental Figure 10. Variant interface, then and now**

Changes to the CIViC Variant representation in the User Interface. The original interface circa 2016[4] (top) had Variants listed to the side of Evidence and Variant Summary, which was moved to accommodate a larger number of Variants and improve searching in the current interface (bottom). **a)** Hover text displays the number of Evidence Items associated with each Variant, pending Suggested Changes to the data associated with the variant and unmoderated (pending) Evidence Items. **b)** Variants can be rapidly filtered by name using the quick filter feature. **c)** Links to the ClinGen Allele Registry[5] are automatically generated when curated Variant Coordinates match an existing Allele Registry Allele. **d)** Curated Aliases can be added to Variant names to enhance variant searching throughout the UI. **e)** CIViC Variant Evidence Scores were added to weight the accumulation of Evidence relative to other CIViC Variants and support the OpenCAP design tool[6].

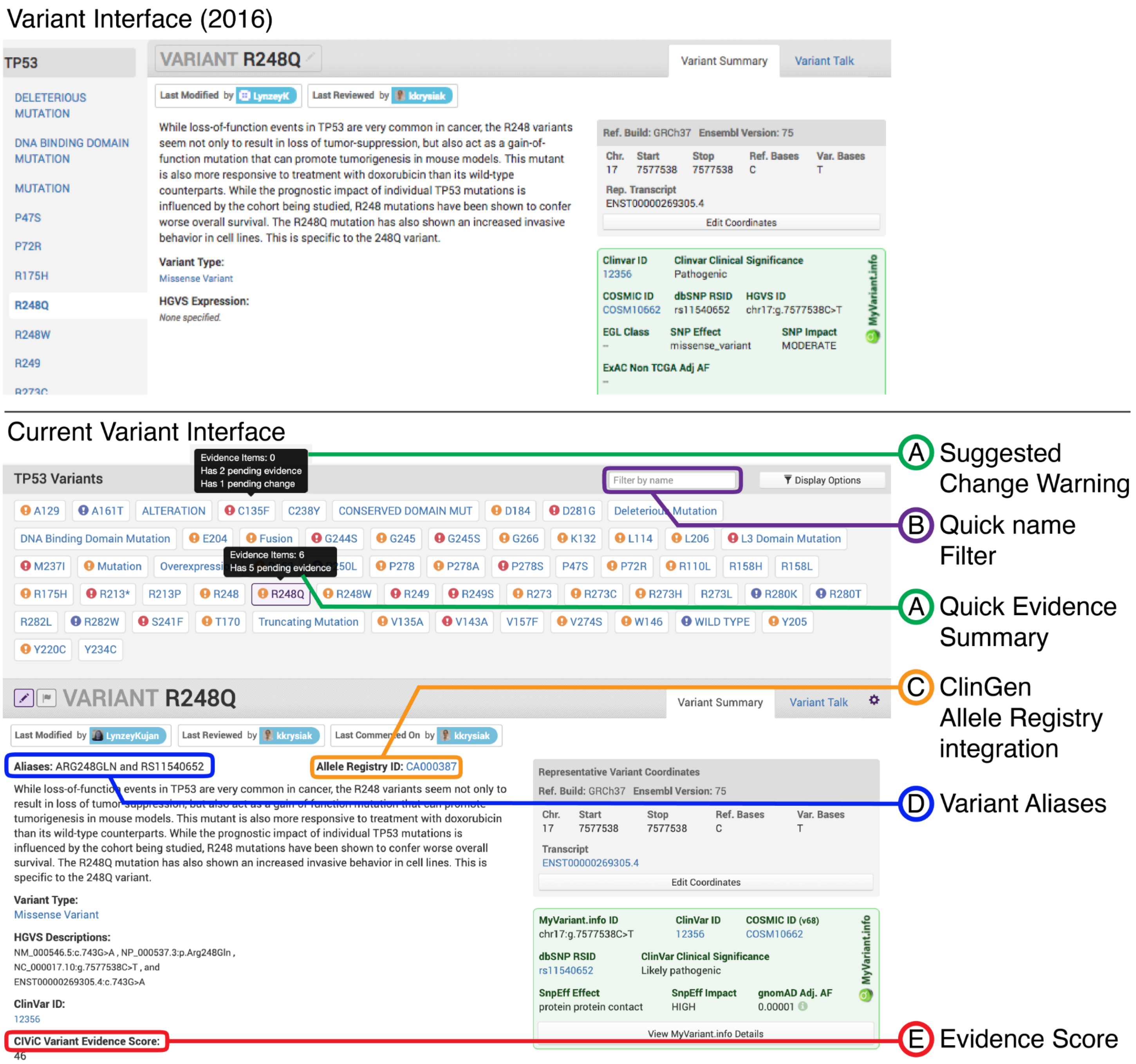

**Supplementary Tables**

**Supplemental Table 1. Peer-reviewed publications associated with CIViC collaborations**

| Project | Organization | Title | PubMed ID |
|---|---|---|---|
| CIViC original publication | WashU | CIViC is a community knowledgebase for expert crowdsourcing the clinical interpretation of variants in cancer[4] | 28138153 |
| Standards for cancer variant interpretation and sharing | Multi-institution initiative / ClinGen Somatic Working Group | ClinGen Cancer Somatic Working Group - standardizing and democratizing access to cancer molecular diagnostic data to drive translational research[2] | 29218886 |
| CIViCmine | Canada's Michael Smith Genome Sciences Centre | Text-mining clinically relevant cancer biomarkers for curation into the CIViC database[7] | 31796060 |
| OpenCAP | WashU | Open-Sourced CIViC Annotation Pipeline to Identify and Annotate Clinically Relevant Variants Using Single-Molecule Molecular Inversion Probes[6] | 31618044 |
| CIViCpy | WashU | CIViCpy: a Python software development and analysis toolkit for the CIViC knowledgebase[8] | 32191543 |
| Virtual Molecular Tumor Board | Multi-institution initiative / GA4GH | Collaborative, Multidisciplinary Evaluation of Cancer Variants Through Virtual Molecular Tumor Boards Informs Local Clinical Practices[9] | 32644817 |
| VICC | Multi-institution initiative / GA4GH | A harmonized meta-knowledgebase of clinical interpretations of somatic genomic variants in cancer[10] | 32246132 |
| MVLD | Multi-institution initiative / ClinGen Somatic Working Group | Adapting crowdsourced clinical cancer curation in CIViC to the ClinGen minimum variant level data community-driven standards[11] | 30311370 |
| CIViC SOP | WashU | Standard operating procedure for curation and clinical interpretation of variants in cancer[12] | 31779674 |
| WikiData | Multi-institution initiative | Wikidata as a knowledge graph for the life sciences[13] | 32180547 |
| DGIdb | WashU | Integration of the Drug-Gene Interaction Database (DGIdb 4.0) with open crowdsource efforts[14] | 33237278 |
